# Clinical evaluation of presepsin considering renal function

**DOI:** 10.1101/604991

**Authors:** Masashi Miyoshi, Yusuke Inoue, Mai Nishioka, Akishige Ikegame, Takayuki Nakao, Koujirou Nagai

**Author notes:** Corresponding author (MM).

## Abstract

Presepsin, a glycoprotein produced during bacterial phagocytosis, has attracted attention as a sepsis marker for bacterial infections. However, since presepsin is affected by renal function, there is a need to separate the evaluation criteria for diagnosis of healthy subjects from that of patients with renal disorder. In this study, we analyzed the influence of kidney function on presepsin concentrations and recalculated the reference range based on the findings. For this purpose, EDTA-whole blood from 47 healthy subjects and 85 patients with chronic kidney disease was collected and used for presepsin measurement by PATHFAST. Presepsin was found to be significantly correlated with creatinine (r = 0.834), eGFRcreat (r = 0.837), cystatin-C (r = 0.845) and eGFRcys (r = 0.879).

Furthermore, in patients with chronic kidney disease at different glomerular filtration rate stages, the presepsin levels showed a significant increasing trend with advancing glomerular filtration rate stage. The reference range, calculated by nonparametric method using a total of 67 cases of healthy volunteers and patients with chronic kidney disease G1, was found to be 59–153 pg/mL, which was notably lower than the standard reference range currently used. Presepsin concentrations were positively correlated with some biomarkers of renal function, indicating that it is necessary to consider the influence of renal function in patients with renal impairment. Further, the recalculated reference range might be more useful for diagnosis and treatment of sepsis than the standard reference currently in use, which likely includes false high values.

## Introduction

Presepsin is a protein whose blood concentrations increase specifically during sepsis. Since its discovery in 2002 in Japan, presepsin has been widely used as a sepsis marker. Membrane-bound CD14, a surface antigen expressed on the cell membrane of monocyte macrophages and granulocytes, is a receptor for bacterial lipopolysaccharide (LPS), which activates cells via TLR4 [1]. In addition, soluble CD14 present in the blood induces the activation of endothelial and epithelial cells without membrane-bound CD14 [2], and plays an important role in sensing invasion of bacteria *in vivo*. Recently, it was reported that granulocyte-mediated bacterial phagocytosis triggers elastase or cathepsin D to proteolytically cleave CD14 to produce presepsin and release it into the blood [3]. Furthermore, it was shown that the concentration of presepsin increases with infection in patients with leucopenia [4], indicating that cells other than monocytes can trigger presepsin production.

Unlike procalcitonin, which has been conventionally used for sepsis diagnosis, presepsin responds very weakly to inflammation such as trauma and burn, and is considered to be highly specific for bacterial infection [5-7]. Compared with conventional markers, presepsin is a good clinical indicator that responds well to changes in the disease state and thus reflects the effect of therapeutics on the condition [8-10].

The presepsin cut-off value for sepsis or infectious disease diagnosis is 400–700 pg/mL (500 pg/mL: Japan) [11]. However, in the blood of a patient with renal disorder such as dialysis patient, the concentration can be higher, because the presepsin is excreted from the kidney [12]. Therefore, it is unsuitable to use the general cut-off values for diagnosis of patients with chronic kidney disease (CKD), as a false high value associated with renal impairment can lead to erroneous judgment.

In the current study, presepsin concentrations were measured in patients with CKD and analyzed for their relationship with the renal function index. In addition, we established a reference range that evaluated the influence of kidney function.

## Materials and methods

### Design and subjects

This study enrolled 85 outpatients with CKD who visited the Tokushima University Hospital from May 2017 to September 2017. All patients were over 18 years of age, and had no history of dialysis and infection. In addition, samples and data were also collected for 47 healthy volunteers without renal dysfunction as controls.

### Blood collection and biochemical analysis

Venous blood was collected after an overnight fasting. Since presepsin concentration increases due to strong agitation, samples after complete blood count (CBC) measurement cannot be used. For this reason, collection was performed with an EDTA-2K blood collection tube exclusively for presepsin measurement. At the time of blood sampling, five gentle inversions were strictly performed.

Estimated glomerular filtration rate (eGFR) of each participant was calculated using the equation provided by the Japanese Society of Nephrology as follows:

eGFRcreat (mL/min/1.73 m^2^) = 194 × creatinine (mg/dL)^-1.094^ × Age^-0.287^ (if female, × 0.739)

eGFRcys (ml/min/1.73 m^2^) = (104 × cystatine [mg/dL]^-1.019^ × 0.996^Age^ [If female, ×0.929]-8)

GFR was categorized according to the KDIGO 2012 by eGFRcys [13].

### Measurement of presepsin

Presepsin concentrations were measured by a compact automated immunoanalyzer “PATHFAST” (LSI Medience Corporation, Tokyo, Japan) based on a chemiluminescent enzyme immunoassay with EDTA-whole blood.

### Ethical approval

This study protocol and consent procedure were approved by the Ethics Committee of Tokushima University Hospital (No. 2699), and performed in compliance with the Helsinki Declaration. Written informed consent was obtained from all patients.

### Statistical analysis

All values are shown as mean ± standard deviation. Results were analyzed as non-parametric variables using Mann-Whitney’s U test for comparison between two groups and using Kruskal-Wallis tests with Bonferroni *post hoc* test for multiple comparisons. Correlation was evaluated by Spearman’s correlation coefficient by a rank test. Statistical analyses were performed with EZR (Saitama Medical Center, Jichi Medical University, Saitama, Japan) [14]. A *P* value < 0.05 was considered significant.

## Results

### Comparison between normal and chronic renal failure

The presepsin level in the chronic renal failure group (CKD G4 · G5) was significantly higher at 410.1 ± 318.9 pg/mL compared with 97.6 ± 27.4 pg/mL in the normal group (*P*= 0.01) (Fig 1).

**Fig. 1.**
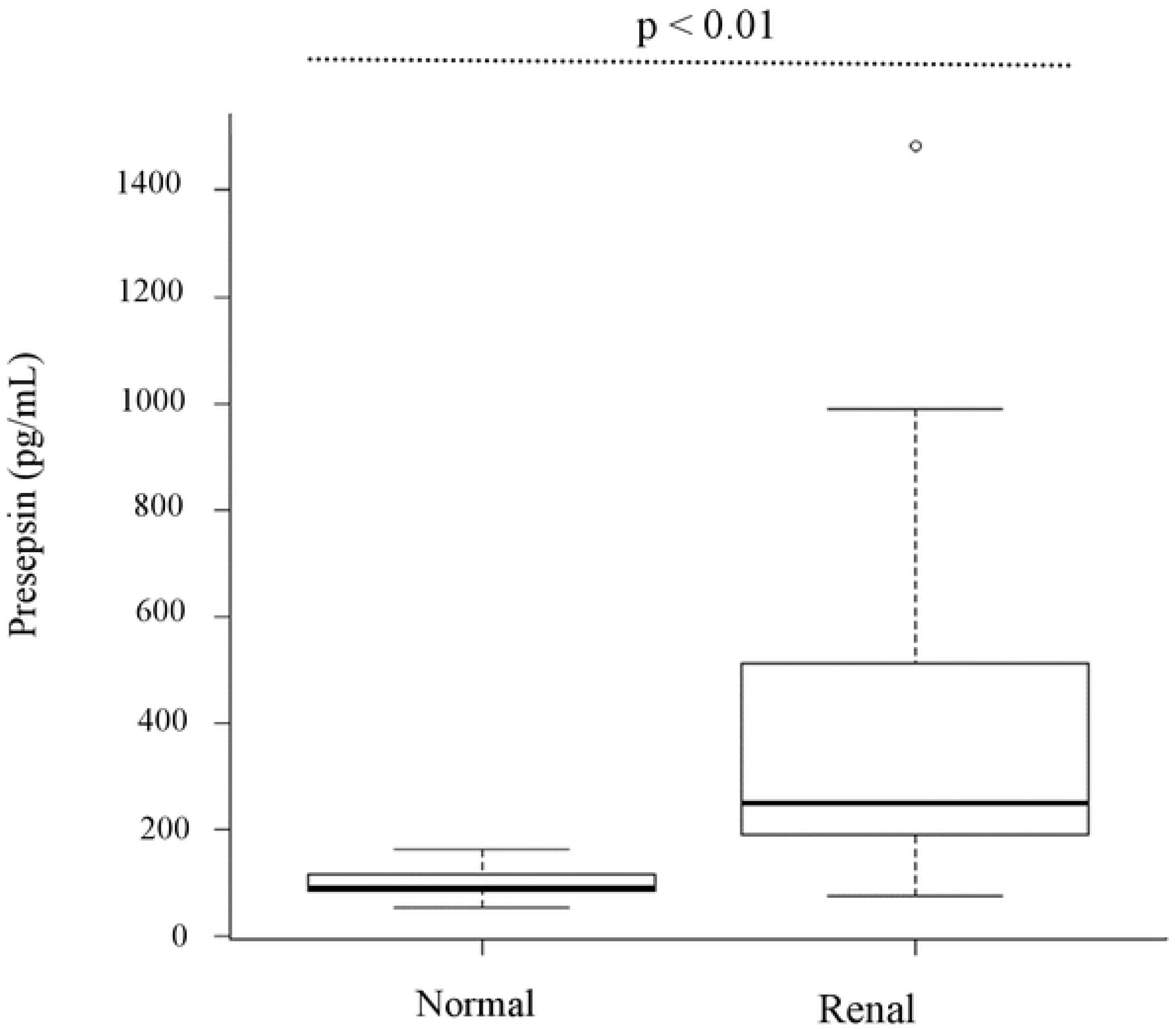
Presepsin levels of control and renal failure group Renal failure group: CKD G4+ G5. *P* value was calculated using the Mann–Whitney U test.

### Correlation with renal functional indices

The presepsin levels were analyzed for their correlation with creatinine, eGFRcreat, cystatin-C, and eGFRcys. The correlation coefficients were as follows: creatinine, r = 0.834; eGFRcreat, r = 0.837; cystatin-C, r = 0.845; and eGFRcys, r = 0.879. All correlations were significant, and the presepsin levels increased with deterioration of renal function (Fig 2).

**Fig. 2.**
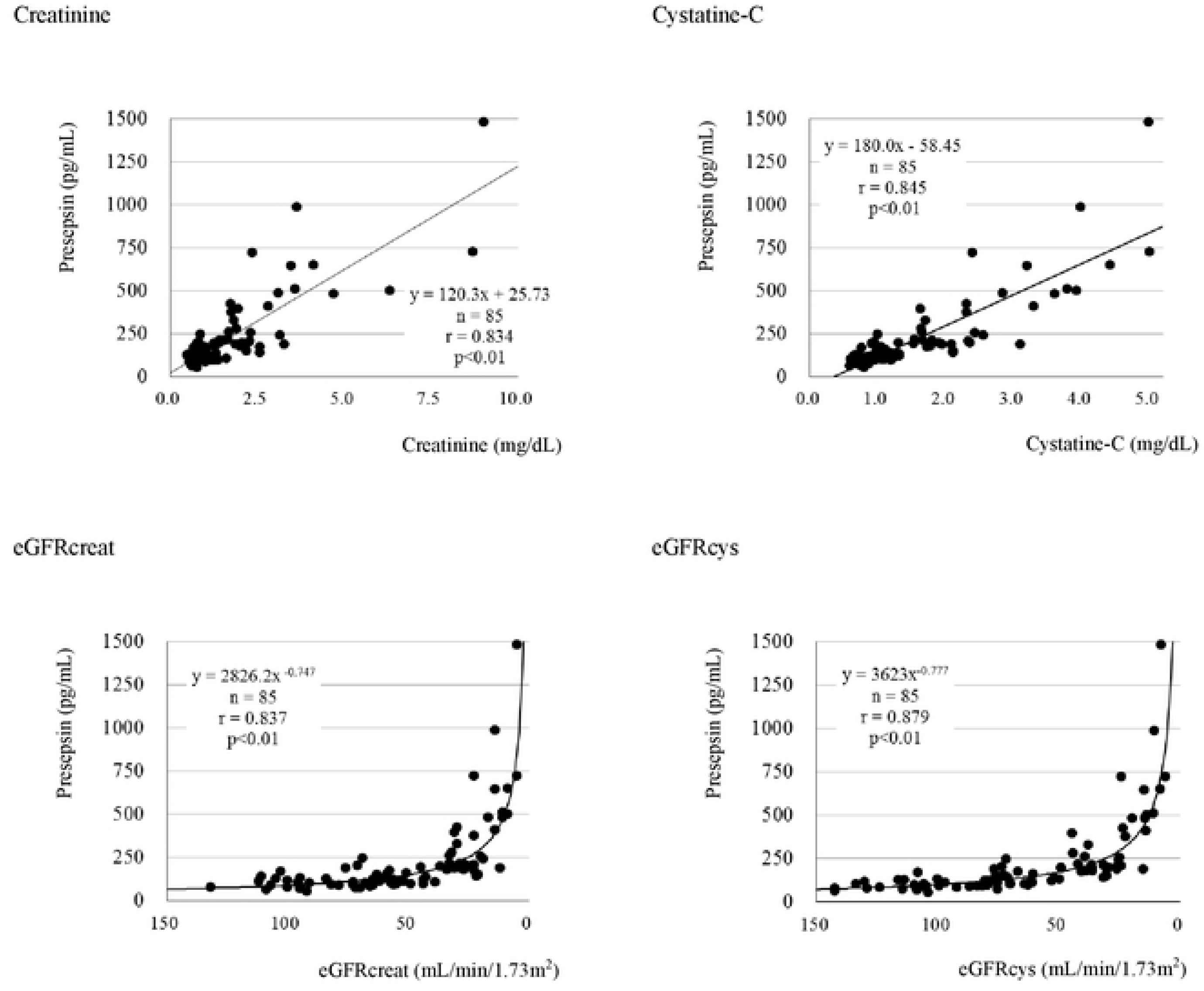
Correlation between presepsin values and renal function index *P* value was calculated with Spearman’s correlation coefficient test. eGFRcreat (mL/min/1.73 m^2^) = 194 × creatinine (mg/dL)^-1.094^ × Age^-0.287^ (if female, × 0.739) eGFRcys (ml/min/1.73 m^2^) = (104 × cystatine [mg/dL]^-1.019^ × 0.996^Age^ [If female, ×0.929]^-8^)

### Stratified comparison in CKD patients

In CKD patients, presepsin levels were stratified by GFR stage classified by eGFRcys (Fig 3). As a result, presepsin levels showed a significant tendency to increase in the CKD G2 patient group, which is regarded as a mildly declining group, as the GFR stage increased. On the other hand, in the CKD G1 patient group, in which renal function was maintained, there was no significant difference between presepsin levels in patients and control individuals.

**Fig. 3.**
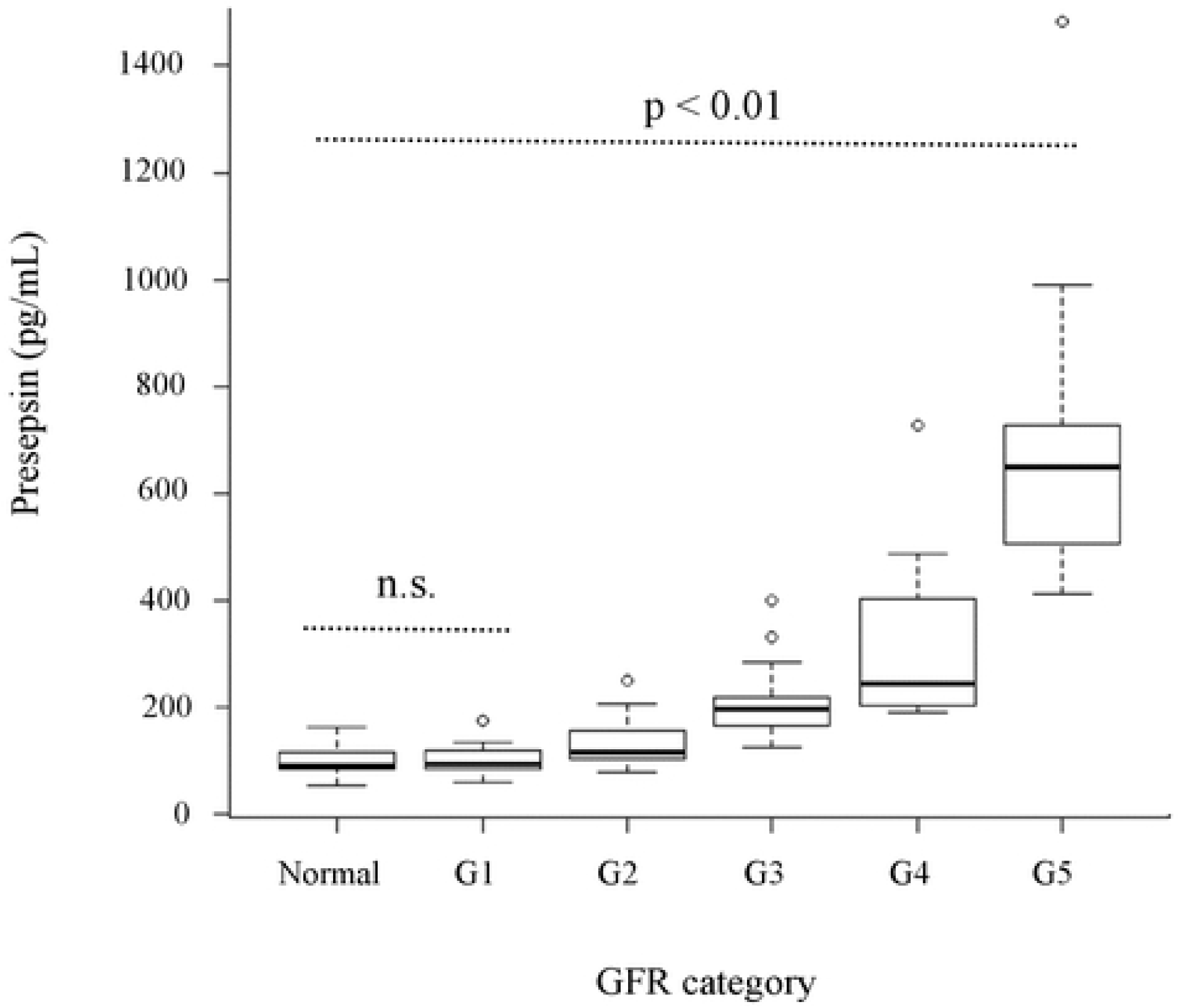
Plot of presepsin concentrations of controls or patients with chronic kidney disease Presepsin levels were stratified by GFR stage classified by eGFRcys. *P* value was calculated using Kruskal-Wallis one-way analysis of variance. *P* value adjustment was calculated using the Bonferroni method.

### Recalculation of reference range considering renal function

A total of 68 cases of the normal group and the CKD G1 patient group were used to determine the reference range calculated by a nonparametric method. The recalculated reference range was 59–153 pg/mL, which was lower than the reference standard range of 98.3–314 pg/mL provided by the reagent manufacturer of PATHFAST.

## Discussion

This study, which excluded the influence of stirring and sampling, clarified that presepsin concentration strongly correlates with various renal function indices and showed a high tendency to increase with renal function deterioration. The molecular weight of presepsin is about 13 kDa, which is almost equivalent to cystatin-C. It is considered that presepsin levels increase and accumulate in blood due to impaired renal function in patients with renal disorders.

Significant correlation was found for all indices examined in this study. Interestingly, eGFRcreat, which is generally used as a renal function index, rather than creatinine, showed the same correlation as creatinine. Since presepsin level was not dependent on the individual’s sex or age, these factors were expected to have no significant influence on the presepsin value. In the stratified comparison, a significant difference was found between eGFRcreat and eGFRcys, although in case of eGFRcys, only the influence from a lower range could be confirmed (S1 Fig). Compared with creatinine, cystatin C was able to reflect the kidney disorder from an earlier stage.

Although presepsin shows an increasing trend from the early stage due to deterioration of renal function, there was no significant difference between healthy subjects and patients in CKD G1 group. Thus, CKD G1 group might be diagnosed as normal. Meanwhile, in patients in CKD G2 group and higher stages, a significant increase in presepsin was observed even in uninfected cases. Therefore, results need to be carefully considered for effect of renal function during sepsis diagnosis in patients with renal impairment.

In addition, false high values of presepsin, arising due to vigorous agitation of the blood samples have been reported, and this could also hinder an accurate clinical assessment. A mechanical stimulus such as agitation leads to formation of a macromolecular complex, which reacts with the anti-presepsin antibody in the reagent in a larger molecular weight fraction. Although the false high value is considered to be due to cross-reaction with this polymer complex, the mechanism of complex formation is not clarified. At present, there is no efficient method other than to avoid agitation of sample at the time of blood collection/delivery, necessitating extreme caution. Although doctors and nurses collect blood on the bedside with caution, fluctuation at the pre-measurement stage can be large, affecting the reliability of the results. Therefore, it is essential to educate clinical practitioners regarding specimen collection.

In this study, recalculation of the reference range using the CKD G1 group and the healthy subject group data indicated remarkably low values that were half of the standard reference range currently in use. The sample group used for calculating the standard reference range did not take into consideration the effects of renal function or agitation, and it is highly probable that these false high values were included. Therefore, it may not be an appropriate sample group. On the other hand, the reference sample group used for calculation in this study excluded false high value cases caused by renal function and agitation, making it a more useful index.

Sepsis diagnosis is not based on only one biomarker, but is a comprehensive diagnosis based on multiple markers and clinical symptoms. The new reference range obtained in this study can be effectively used for early diagnosis of sepsis. For example, mild sepsis can be positively identified with elevated levels of presepsin in the range of 300–500 pg/mL.

In conclusion, by excluding the effect of stirring during sampling, our study showed that presepsin levels exhibit an increasing trend as GFR decreases. Currently, the false high values in the standard reference range are largely ignored. Therefore, the use of the newly recalculated reference range, which excludes the effects of renal function and agitation, is expected to improve efficiency in sepsis diagnosis, particularly at the early stage.

## Acknowledgement

We would like to thank Naofumi Yoda for her helpful insights. We would also like to thank Editage for English language editing.

---

**Table 1.** Clinical characteristics of participants in the different GFR categories Data are presented as mean ± standard deviation, as number (%). GFR categories: Categorized according to the KDIGO 2012 by eGFRcys. G1: eGFRcys ≥ 90 ml/min/1.73 m^2^; G2: eGFRcys = 60–90 ml/min/1.73 m^2^; G3: eGFRcys = 30–60 ml/min/1.73 m^2^; G4: eGFRcys = 15–30 ml/min/1.73 m^2^; G5: eGFRcys ≤ 15 ml/min/1.73 m^2^.

|  | Normal | G1 | G2 | G3 | G4 | G5 |
| --- | --- | --- | --- | --- | --- | --- |
| N | 47 | 20 | 26 | 19 | 11 | 9 |
| age | 30.8 ± 10.5 | 40.6 ± 11.2 | 56.7 ± 12.5 | 61.8 ± 14.9 | 66.0 ± 13.3 | 66.9 ± 13.6 |
| male,n (%) | 21 (44.7) | 10 (50.0) | 20 (76.9) | 15 (78.9) | 6 (54.5) | 6 (66.6) |
| AST (U/L) | 18.0 ± 6.2 | 23.2 ± 12.7 | 22.9 ± 8.6 | 24.2 ± 8.0 | 19.1 ± 6.0 | 21.4 ± 7.6 |
| ALT (U/L) | 16.1 ± 15.5 | 23.8 ± 12.1 | 19.2 ± 9.1 | 20.6 ± 11.1 | 11.7 ± 3.7 | 15.2 ± 4.1 |
| creatinine (mg/dL) | 0.718 ± 0.15 | 0.684 ± 0.16 | 0.952 ± 0.22 | 1.716 ± 0.45 | 2.245 ± 0.70 | 5.143 ± 2.33 |
| cystatine-C (mg/dL) | no data | 0.710 ± 0.07 | 1.007 ± 0.11 | 1.657 ± 0.24 | 2.415 ± 0.33 | 4.021 ± 0.66 |
| urea nitrogen (mg/dL) | 13.2 ± 3.2 | 12.4 ± 3.1 | 15.2 ± 4.1 | 26.4 ± 6.9 | 36.8 ± 8.0 | 56.7 ± 20.0 |
| eGFR creat (ml/min/1.73m<sup>2</sup>) | 93.0 ± 12.9 | 92.4 ± 18.6 | 63.3 ± 14.3 | 33.0 ± 9.9 | 23.2 ± 6.2 | 10.2 ± 3.6 |
| eGFR cys (ml/min/1.73m<sup>2</sup>) | no data | 114.2 ± 14.8 | 73.7 ± 7.2 | 40.1 ± 7.4 | 23.9 ± 3.7 | 11.3 ± 3.3 |
| white blood cell (*10<sup>9</sup>/L) | 5.5 ± 1.2 | 6.4 ± 2.3 | 6.1 ± 1.6 | 5.5 ± 1.4 | 5.8 ± 1.8 | 6.5 ± 2.1 |
| hemoglobin (g/dL) | 13.6 ± 1.5 | 14.4 ± 1.6 | 13.8 ± 1.5 | 13.1 ± 1.6 | 11.7 ± 1.3 | 10.8 ± 1.4 |
| hematcrit (%) | 41.0 ± 4.0 | 42.6 ± 4.1 | 40.7 ± 4.0 | 38.4 ± 4.6 | 35.6 ± 3.5 | 33.1 ± 3.4 |
| platelets (*10<sup>9</sup>/L) | 247.1 ± 49.4 | 242.1 ± 64.6 | 245.4 ± 52.1 | 215.3 ± 43.7 | 353.6 ± 463.9 | 188.2 ± 43.5 |

**S1 Fig:**
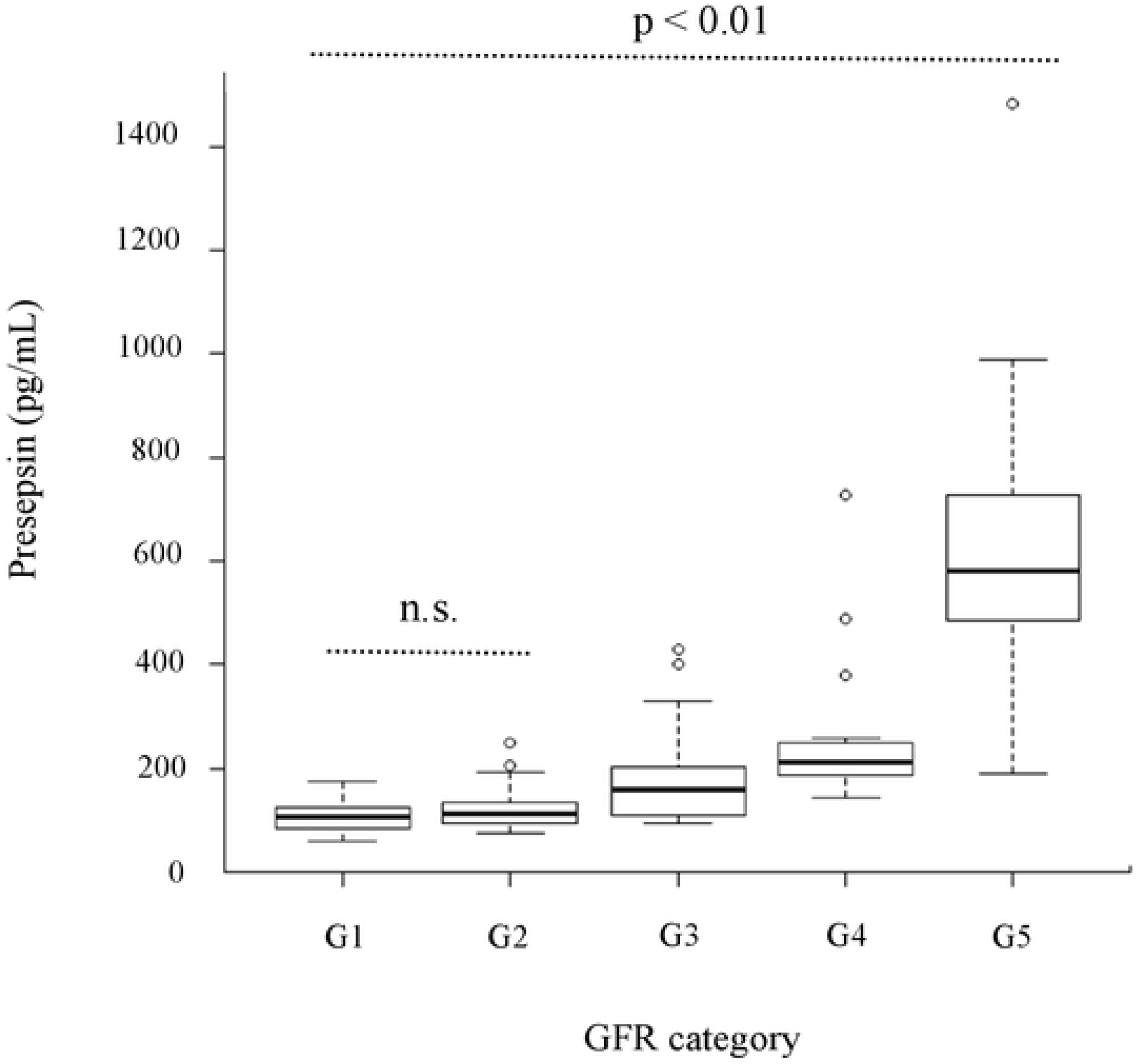
Plot of presepsin concentrations of patients with chronic kidney disease Presepsin levels were stratified by GFR stage classified by eGFRcreat. *P* value was calculated using Kruskal-Wallis one-way analysis of variance. *P* value adjustment was calculated using the Bonferroni method.

